# Investigating EMT-mediated resistance to EGFR tyrosine kinase inhibitors in NSCLC using innovative organoid models

**DOI:** 10.1101/2024.02.08.579426

**Authors:** Nobuaki Kobayashi, Seigo Katakura, Nobuhiko Fukuda, Shuhei Teranishi, Sousuke Kubo, Chisato Kamimaki, Hiromi Matsumoto, Kohei Somekawa, Ayami Kaneko, Yoshihiro Ishikawa, Koji Okudela, Keisuke Sekine, Takeshi Kaneko

## Abstract

**Background:** Epithelial-mesenchymal transition (EMT) has emerged as a key mechanism underlying resistance to epidermal growth factor receptor (EGFR) tyrosine kinase inhibitors (TKIs) in EGFR-mutant non-small cell lung cancer (NSCLC). However, the intricacies of EMT-mediated resistance, driven by tumor microenvironment (TME) interactions, remain enigmatic. This study aimed to probe EMT-induced resistance in NSCLC using innovative in vitro organoid models.

**Methods:** We generated organoids by co-culturing EGFR-mutant NSCLC cells (HCC827, H1975), mesenchymal stem cells and endothelial cells. Drug susceptibility was compared between organoids and spheroids (cancer cells only) using EGFR TKIs - Gefitinib, Afatinib, Osimertinib. EMT marker (E-cadherin, ZEB1) expression was analyzed via immunofluorescence and western blotting. The effects of Bevacizumab and miR200c on overcoming resistance were also investigated.

**Results:** The study identified a significant link between EMT and EGFR-TKI resistance. Notable findings included the decrease of E-cadherin and an increase in ZEB1, both of which influenced EMT and resistance to treatment. Bevacizumab showed promise in improving drug resistance and mitigating EMT, suggesting an involvement of the VEGF cascade. Transfection with miR200c was associated with improved EMT and drug resistance, further highlighting the role of EMT in TKI resistance.

**Conclusions:** This study offered vital insights into EMT-driven EGFR TKI resistance, highlighting the utility of organoid models in evaluating resistance modulated by TME interactions. Our findings reveal promising directions for overcoming EMT-mediated resistance involving Bevacizumab and miR200c, warranting further in vivo validation.

## Introduction

Non-small cell lung cancer (NSCLC) remains the foremost cause of cancer-related deaths worldwide[1]. The identification of oncogenic driver mutations has transformed the therapeutic landscape for NSCLC, with EGFR mutations observed in approximately 15-50% of advanced cases[2]. Predominantly comprising exon 19 deletions or L858R mutations in exon 21, these EGFR alterations serve as key targets for EGFR tyrosine kinase inhibitors (TKIs) - a therapeutic intervention that significantly enhances the prognosis of inoperable NSCLC patients[3], [4]. Nevertheless, acquired resistance to EGFR-TKIs remains a considerable clinical challenge, typically emerging between 11 to 18 months following treatment initiation[5], [6]. Several mechanisms, including genetic mutations[7], [8], histological transformation[9], and the epithelial-mesenchymal transition (EMT)[10], contribute to this resistance, with EMT particularly instrumental in enabling cancer cell invasion and metastasis. Accordingly, inhibiting EMT emerges as a promising strategy to surmount drug resistance.

The complexity of these tumor microenvironment (TME)-dependent resistance mechanisms necessitates sophisticated in vitro models for their elucidation. Among these, organoids - self-organizing, multicellular structures capable of mimicking in vivo physiology and tissue-specific functions - hold substantial promise. Through the co-culture of lung cancer cells, human mesenchymal stem cells (MSCs), and human umbilical vein endothelial cells (HUVECs) on low attachment plates, organoids that encapsulate the intricate cellular interactions integral to EMT can be generated[11]– [13].

In this study, we constructed an in vitro EMT model of EGFR-mutant lung cancer organoids to identify potential EMT-regulating compounds. We delved into the interplay between the TME and EGFR-TKI resistance, aiming to contribute to the broader understanding of EMT and its role in anticancer drug resistance.

## Materials and Methods

### Cell lines

The luciferase-tagged human lung adenocarcinoma cell lines HCC827 and H1975 were procured from the Japanese Cancer Research Resources Bank. HCC827 cell lines exhibited Exon 19 deletions, while H1975 had L858R and T790M mutations on the EGFR gene. The cancer cells were maintained in Roswell Park Memorial Institute (RPMI) medium (Sigma-Aldrich, St. Louis, MO, USA), supplemented with 10% fetal bovine serum (FBS) (Biowest, Nuaillé, France) and 1% Pen/Strep (Gibco, Waltham, MA, USA) in a 5% CO_2_ environment at 37℃. MSC and HUVEC were procured from TAKARA (Shiga, Japan). MSCs were cultivated in Mesenchymal Stem Cell Growth Medium 2 (PromoCell, Heidelberg, Germany) at 37℃ in 5% CO2. HUVECs were cultivated in Endothelial Cell Growth Medium 2 (EGM) (PromoCell) at 37℃ in 5% CO_2_. Both MSCs and HUVECs were utilized before their 10th passage.

### The construction of human lung cancer spheroids and organoids

Matrigel (Corning, Corning, NY, USA) was combined in equal parts with RPMI medium. This mixture (30 µL) was seeded onto a cold 96 well plate (Corning), which had been cooled using phosphate-buffered saline (PBS) (Nihon Gene, Tokyo, Japan). Following an incubation period at 37℃ for 1 hour, 1.11×10^5^ cancer cells were seeded to create spheroids. For organoids, 3.0×10^4^ cancer cells, 6.0×10^4^ MSCs, and 2.1×10^4^ HUVECs were seeded. The resultant cell pellets were resuspended in 30 µL drops on a 96 well plate. After a 30-minute incubation at 37℃, 200 µL of culture medium (a 1:1 mixture of RPMI medium and EGM) was added to each well upon complete gelation. Plates were then transferred to humidified 37℃/ 5% CO_2_ incubators to allow 3D structures to form for subsequent analyses.

### Mimicking micro RNA200c (miR200c)

Lipofectamine RNAiMAX Reagent (Invitrogen, Carlsbad, CA, USA) and mimic miR200c (Eurofins Genomics, Tokyo, Japan) were directly added to the culture medium for organoid transfection. S-TuD-NC (Eurofins Genomics) was used as a negative control. Each well received 1 pmol of final mimic miR200c. The sequence was as follows: mimic miR200c (upstream primer, 5’-TAATACTGCCGGGTAATGATGGA-3’). The transfection incubation period was one day.

### Total RNA isolation, reverse-transcription and polymerase chain reaction (PCR)

Total RNA was extracted from the organoids using the RNeasy mini kit (QIAGEN, Venlo, Netherlands). The concentration of the extracted total RNA was determined using the Nanodrop 2000 system (Thermo Fisher Scientific, Waltham, MA, USA). The Mir-X miRNA First-Strand Synthesis Kit (TAKARA, Shiga, Japan) was used to convert RNAs into complementary DNA (cDNA). Reverse-transcription-quantitative PCR (RT-qPCR) was conducted using the Mir-X miRNA qRT-PCR TB Green Kit (TAKARA), the reverse-transcription samples, and the miR200c primers (upstream primer, 5’- TAATACTGCCGGGTAATGATGGA-3’) (Eurofins Genomics) on the CFX96 Touch Real-Time PCR Detection System (Bio-Rad, Hercules, CA, USA). U6 primers (TAKARA) were used to normalize miRNA expression via the 2^-ΔΔCq^ method (ΔCq = Cq_target_ − Cq_rereference_).

### Immunohistochemical analyses

The spheroids and organoids were cultured for 72 hours, fixed with 4% paraformaldehyde overnight, then paraffin-embedded and sectioned. After deparaffinization, morphology was examined via Hematoxylin and Eosin (HE) staining. These slices were stained with anti-EGFR antibody (Nichirei Biosciences, Tokyo, Japan) and anti-mouse IgG conjugated with horseradish peroxidase (HRP) (Dako, Carpinteria, CA, USA) for immunohistochemical staining.

### Cell viability

After 24 hours from seeding, 1 nM-10 µM of EGFR-TKIs (Gefitinib, Med Chem Express, NJ, USA / Afatinib, Med Chem Express / Osimertinib, Med Chem Express) were added. The spheroids and organoids were co-cultured with EGFR-TKIs ± Bev for 72 hours. The 100 µL of cell culture lysis reagent (Promega, WI, USA) was added and suspended. The mixture was frozen and thawed repeatedly. After 24 hours, the 20 µL of samples with 20 µL of luciferase substrate (Promega) were transferred to the 96 well plate (Thermo Fisher Scientific, MA, USA) and luciferase reporter activities were measured by luminometer (PerkinElmer, MA, USA) to calculate the survival rate of cells. Averages of IC_50_ from at least three independent experiments.

### Immunofluorescence analyses

Preparation of samples for immunofluorescence analyses paralleled the procedure used for immunohistochemical studies. Sections were incubated overnight at 4℃ with anti-E-cadherin antibody (BD Biosciences, NJ, USA) and anti-ZEB1 antibody (Novus Biologicals, CO, USA). Following this, the sections were treated with goat-anti-mouse IgG Alexa Fluor 488 and goat-anti-mouse IgG Alexa Fluor 555 for 1 hour (both from Thermo Fisher Scientific, MA, USA). The staining procedure was conducted using the Antifade Reagent with DAPI (Cell Signaling Technology, MA, USA), and cellular observation was performed under a BZ-X800 fluorescence microscope (Keyence Corporation, Osaka, Japan). This procedure was also followed for spheroids and organoids treated with Bevacizumab.

### Western blotting

Total protein was extracted from spheroids and organoids using lysis buffer supplemented with a protease/phosphatase inhibitor cocktail (both from Cell Signaling Technology, MA, USA). Protein concentrations were measured using a Nanodrop 2000 system (Thermo Fisher Scientific, MA, USA). Proteins were separated by 10% sodium dodecyl sulfate-polyacrylamide gel electrophoresis (Bio-Rad Laboratories, CA, USA) and transferred onto polyvinylidene difluoride membranes (Invitrogen, CA, USA). Membranes were blocked with 3% skimmed milk for 1 hour, then incubated with anti-E-cadherin antibody (BD Biosciences, NJ, USA), anti-ZEB1 antibody (Novus Biologicals, CO, USA), and anti-β-actin antibody (Cell Signaling Technology, MA, USA) at 4℃ overnight. Membranes were subsequently reacted with horseradish peroxidase (HRP)- linked secondary antibodies for 1 hour at room temperature. Band intensities were quantified using an ImageQuant LAS 500 system (GE Healthcare, IL, USA). The same experiments were repeated for spheroids and organoids treated with Bevacizumab or transfected with miR200c. Protein quantification was conducted using ImageJ (National Institutes of Health, MD, USA).

### Statistical analysis

Data representation and statistical analysis were conducted using JMP 15.0.0 software program (SAS Institute, Cary, NC, USA). The Mann-Whitney U test was employed to assess differences between groups. Unless otherwise stated, data were presented as the mean ± standard error of the mean. Significance was considered at p<0.05.

## Results

### In Vitro Development of EGFR-Mutant Lung Cancer Organoids

In an attempt to construct an innovative in vitro model, EGFR-mutant lung cancer organoids were established using HCC827 and H1975 cells, which are known to carry EGFR mutations. Notably, these cell lines were successful in forming complex, three-dimensional lung cancer organoids when co-cultured with MSCs and HUVECs.

Detailed organoid formation was demonstrated in Figure 1, featuring immunohistochemical staining. Specifically, positive EGFR staining provided a distinct demarcation between the cancer cells and the stromal components, corroborating the successful incorporation of these cell types into the organoid structure. It was further noted that, compared to spheroids, the organoids showed a higher degree of stromal tissue composition, hinting at their superior potential in mimicking the natural tumoral microenvironment in vivo. Intriguingly, the morphological differences between the two EGFR-mutant lung cancer cell lines, HCC827 (Figure 1A) and H1975 (Figure 1B), remained insignificant within the organoid context. This consistent morphology suggests a potential underlying uniformity in the behavior of EGFR-mutant cells within organoids.

**Figure 1:**
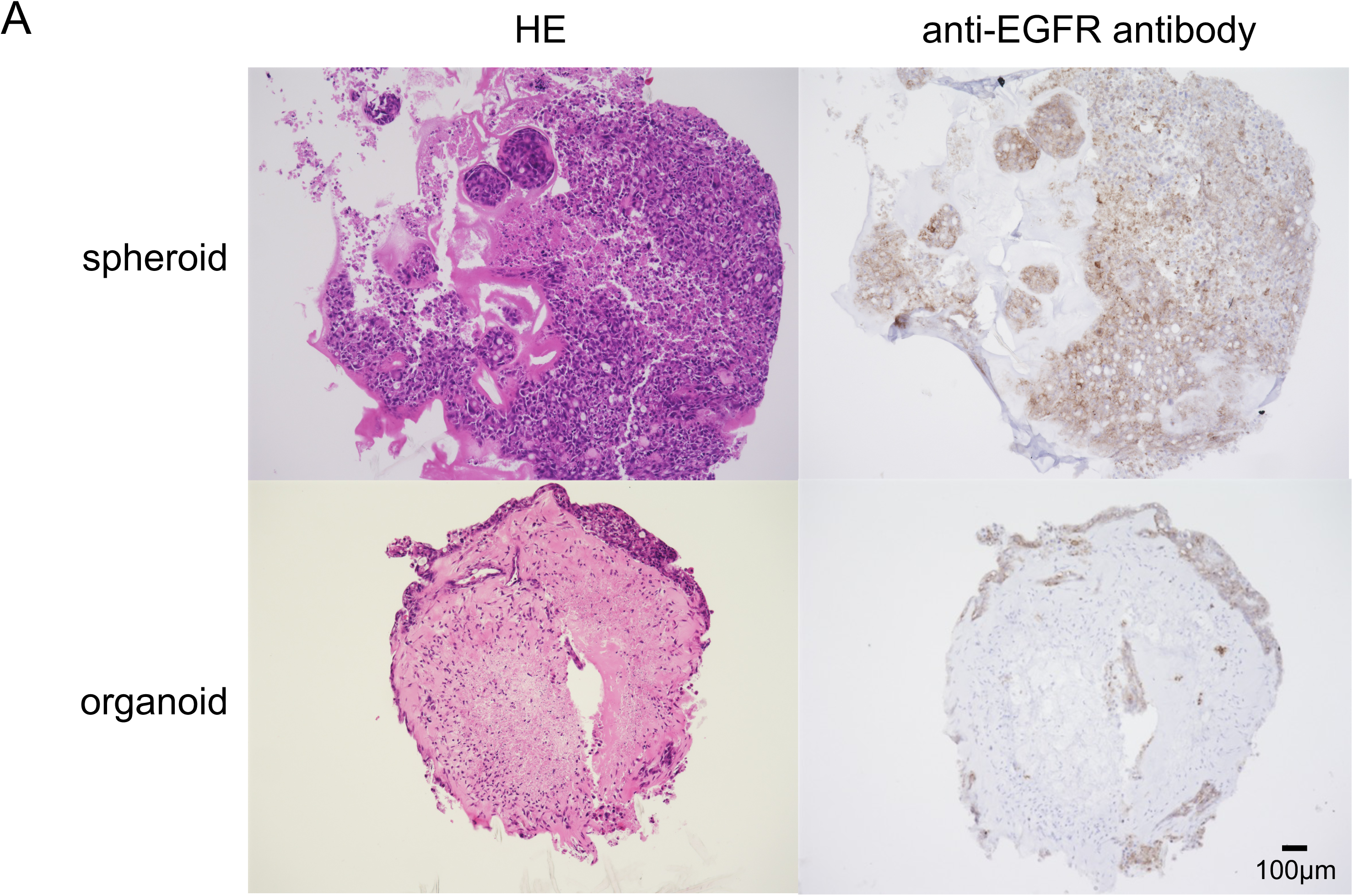
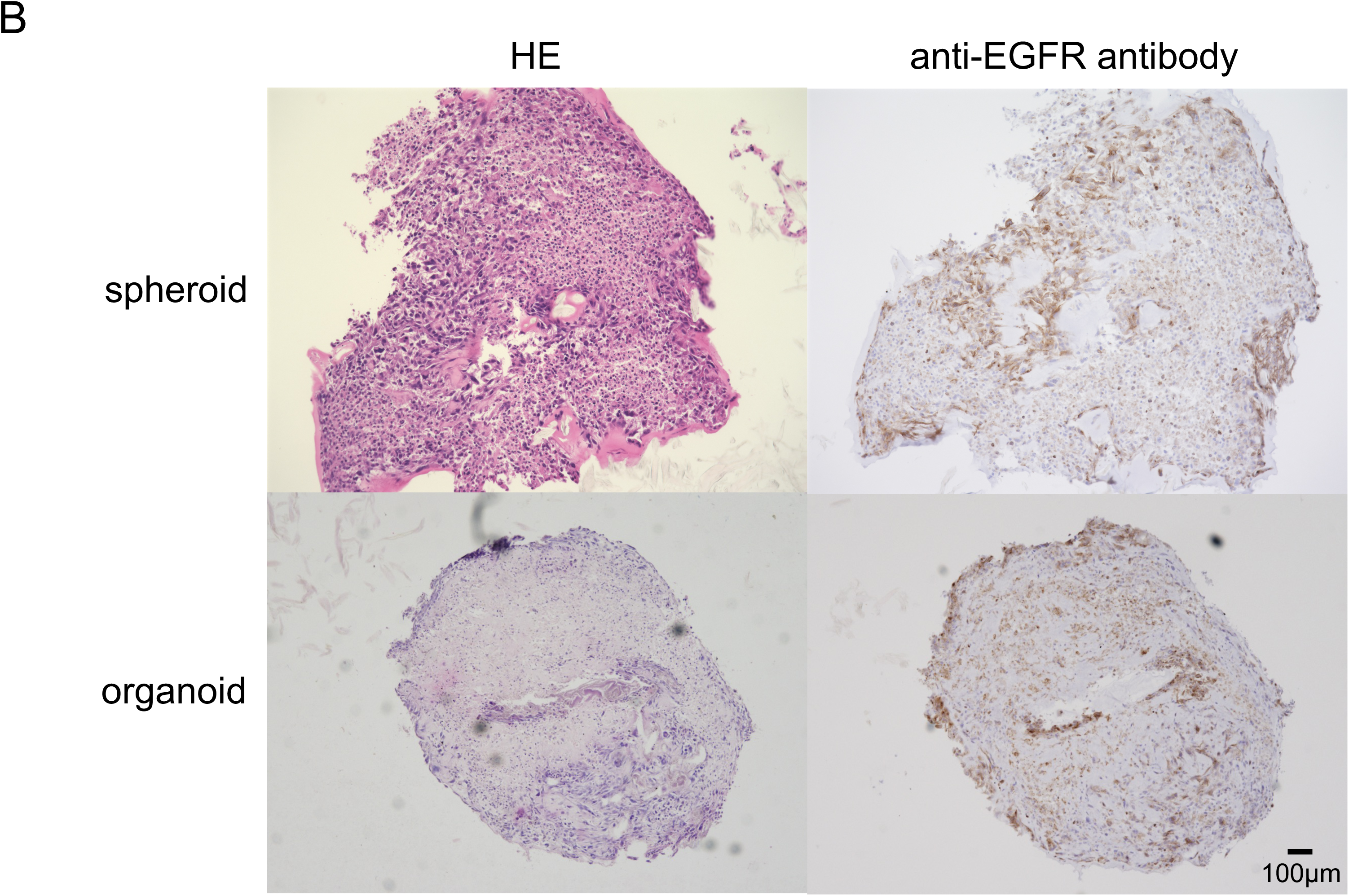
Construction and microscopic findings of spheroids and organoids using EGFR-mutant lung cancer cells. (A) Hematoxylin and eosin (HE) staining and EGFR immunostaining of HCC827-derived spheroids and organoids. (B) HE and EGFR staining of structures generated from H1975 cells.

### Differential Susceptibility of Spheroids and Organoids to EGFR-TKI

To elucidate the differential response of spheroids and organoids to EGFR-TKI therapy, a series of experiments involving luciferase assays were conducted. The ensuing data, presented in Figure 2, unraveled marked differences in the therapeutic susceptibility of the two models.

**Figure 2:**
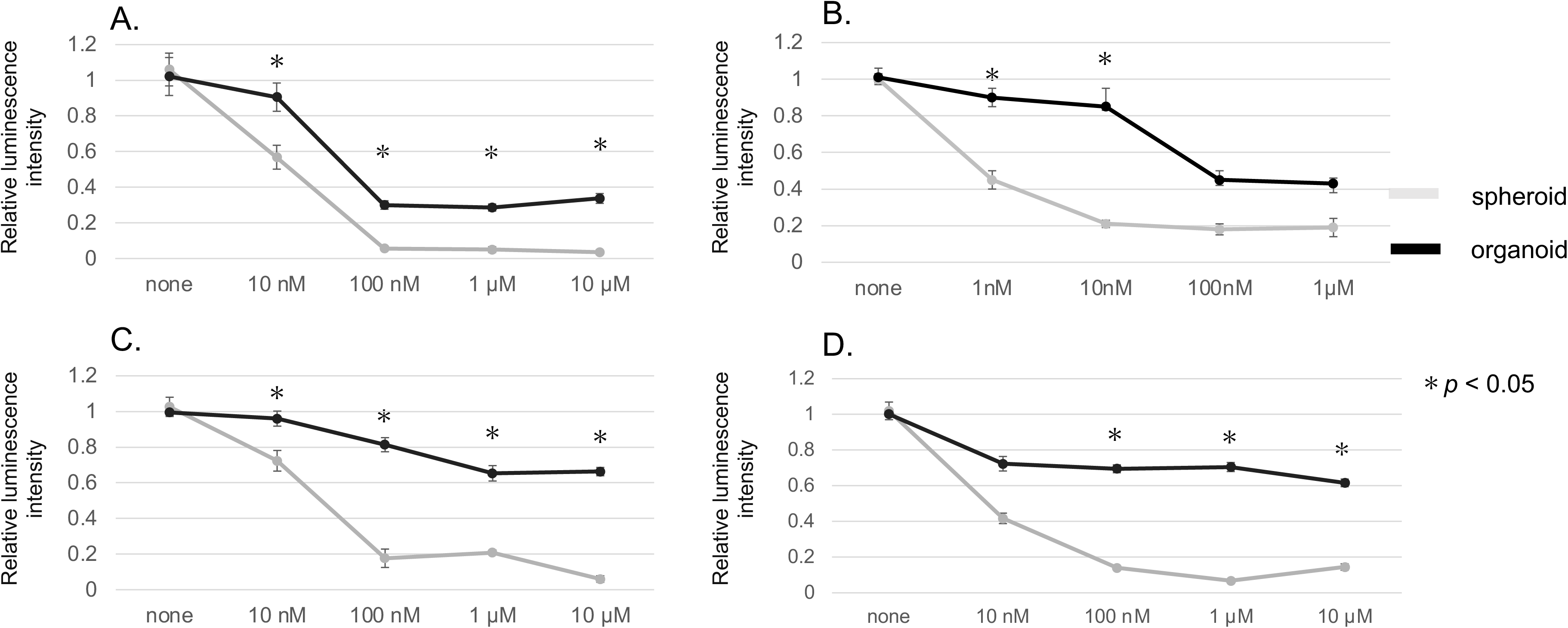
Assessment of cell viability in response to various EGFR-TKIs in spheroids and organoids derived from cancer cells, determined by a luciferase assay. Each panel displays the influence of an individual EGFR-TKI on spheroids or organoids: (A) Gefitinib on HCC827, (B) Afatinib on HCC827, (C) Osimertinib on HCC827, and (D) Osimertinib on H1975. Each experiment was repeated three times (n=3), with *p<0.05 (determined by the Mann-Whitney U test) indicating a statistically significant difference in cell viability.

In Figure 2A, the cell viability of HCC827 cells, subsequent to incubation with Gefitinib, is shown. Figures 2B and 2C depict the respective viability of HCC827 cells following treatment with Afatinib and Osimertinib. Figure 2D presents the results of the viability of H1975 cells post-treatment with Osimertinib.

A prominent finding from the experiments was the considerable elevation in IC50 values for all employed EGFR-TKIs in organoids in contrast to their spheroid equivalents (Figure 2A-D). This observation implies a heightened resistance to EGFR-TKI therapy within the organoid model.

### Epithelial-to-Mesenchymal Transition in Organoids: Immunofluorescence Analysis of EMT Markers

To substantiate the hypothesis of EMT-induced resistance to EGFR-TKI in organoids, immunofluorescence analyses were carried out to assess the expression of key EMT markers. The findings were visually represented in Figure 3, indicating significant variations between organoids and spheroids.

**Figure 3:**
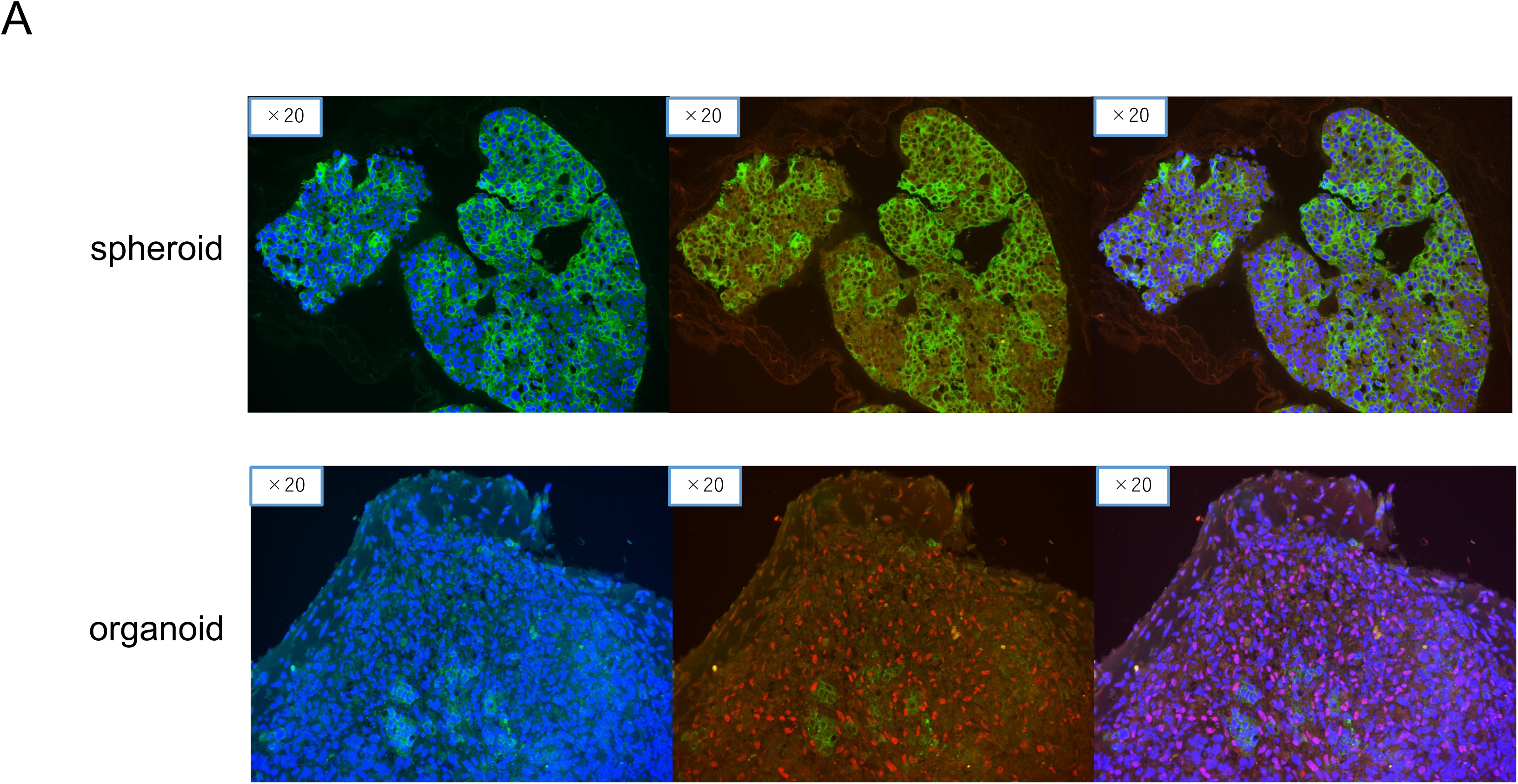
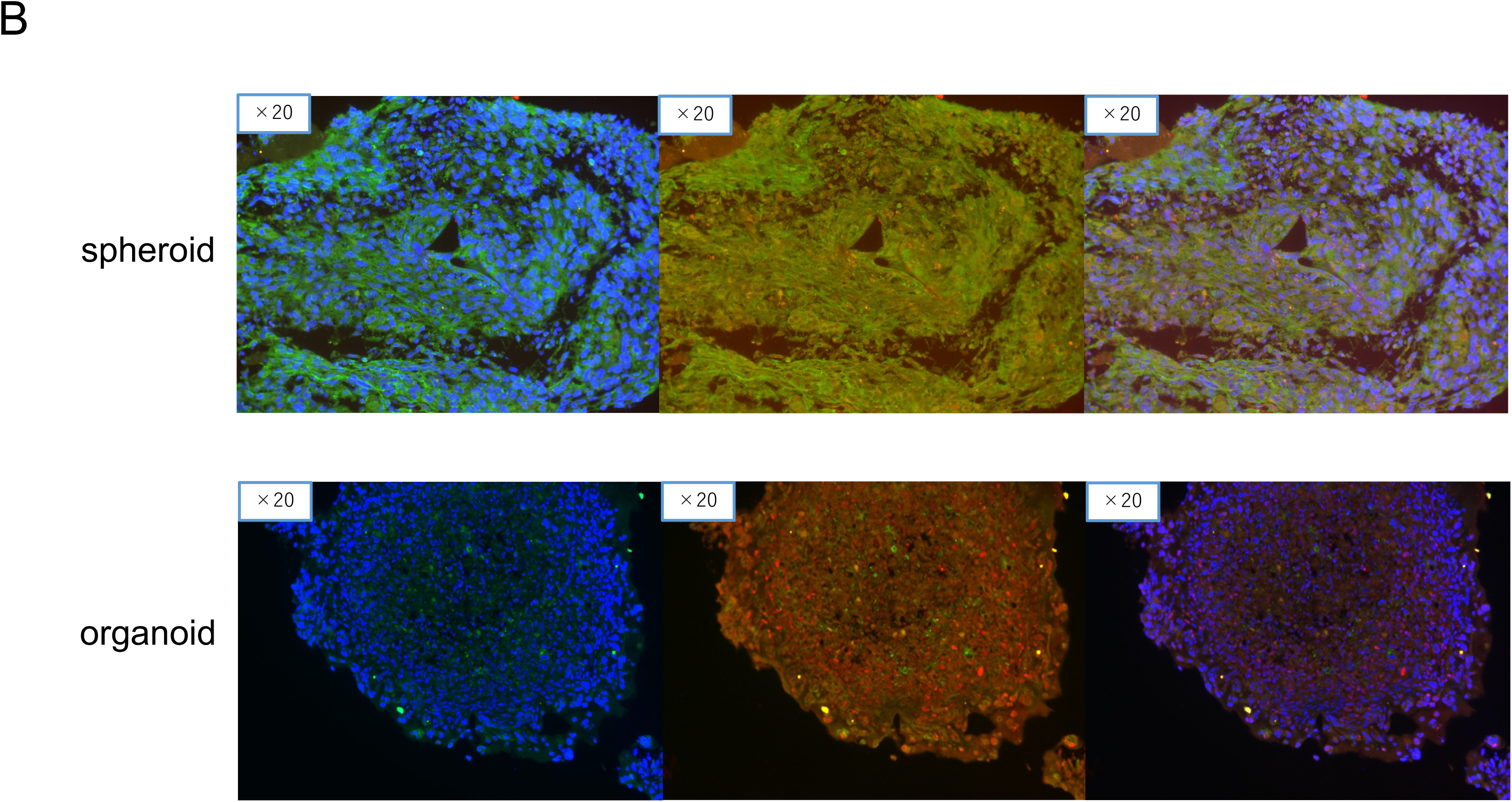
Immunofluorescence imaging of Epithelial Cadherin (E-cadherin) and Zinc-finger-enhancer Binding Protein 1 (ZEB-1) in spheroids and organoids derived from lung cancer cell lines. The images illustrate cellular staining patterns for DAPI (blue, indicating cell nuclei), E-cadherin (green, an epithelial marker), and ZEB-1 (red, a mesenchymal marker). (A) Visualization of these markers in HCC827-derived structures, and (B) the same in H1975-derived structures.

Figure 3A illustrated the upregulation of Epithelial Cadherin (E-cadherin, an epithelial marker) and the concurrent downregulation of Zinc-finger-enhancer binding protein 1 (ZEB-1, a marker for EMT) in HCC827 organoids as compared to HCC827 spheroids. This expression pattern aligns with the characteristic signature of EMT, suggesting a more pronounced induction of EMT in organoids than in spheroids.

An analogous observation was made in the case of H1975 cells, as depicted in Figure 3B. The data confirmed a similar trend of EMT marker expression, reinforcing the suggestion that EMT may play a pivotal role in the observed EGFR-TKI resistance in organoids.

### Modulation of EMT and EGFR-TKI Resistance in Organoids by Bevacizumab

The potential influence of Bevacizumab (Bev), an antibody against vascular endothelial growth factor (VEGF), on EMT and EGFR-TKI resistance in organoids was explored in the next.

A luciferase assay was employed to discern the combined effects of Bev and EGFR-TKI on EGFR-mutant cancer cells. As demonstrated in Figure 4A for HCC827 cells and in Figure 4B for H1975 cells, the IC50 of the combined Osimertinib and Bev treatment was significantly lower than that of Osimertinib alone. These findings suggest an improvement in therapeutic efficacy, possibly linked to EMT suppression, with the addition of Bev.

**Figure 4:**
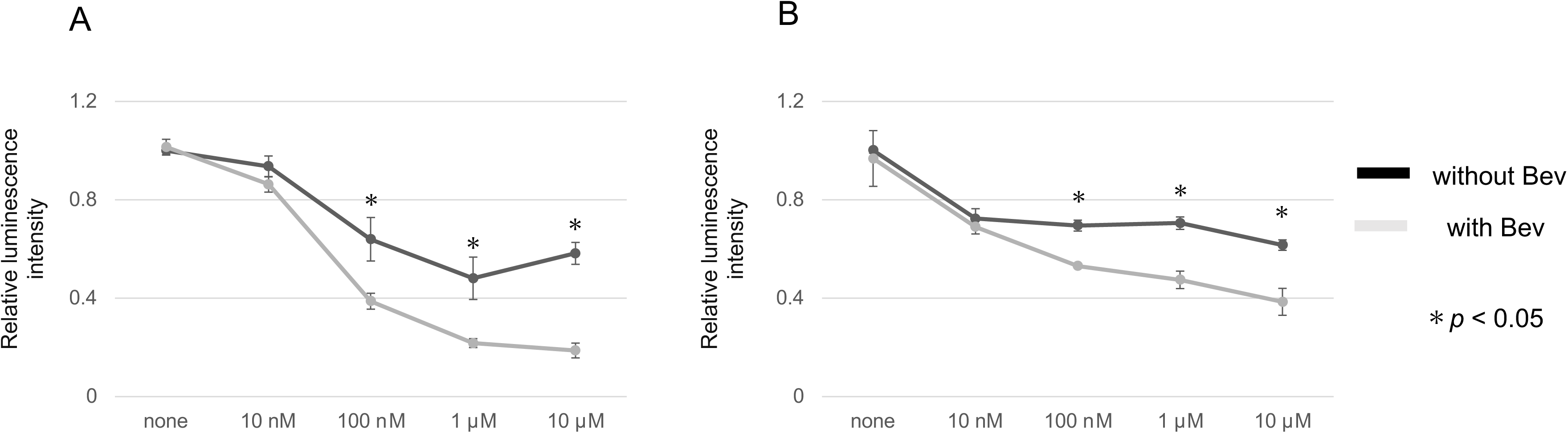
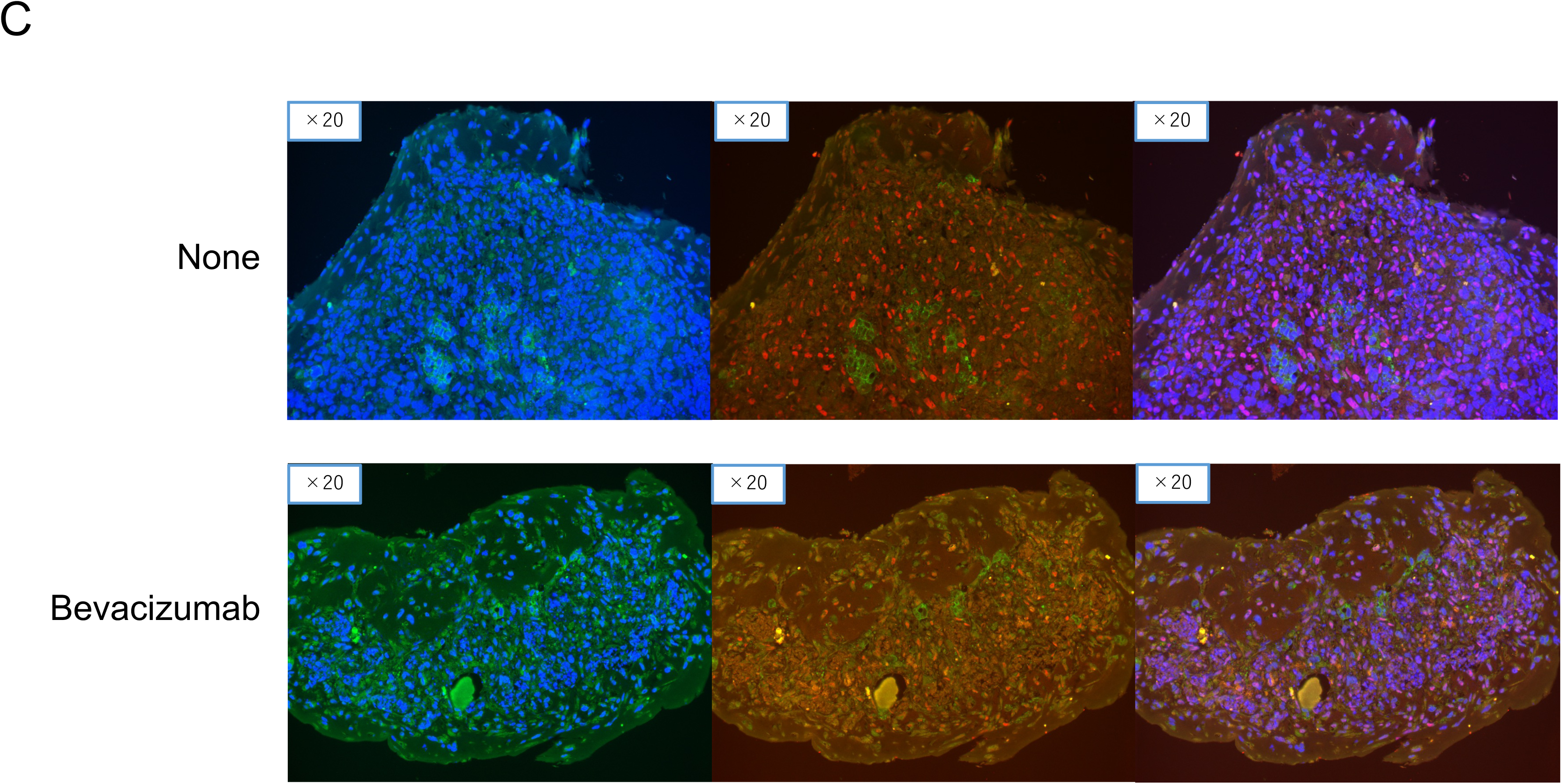
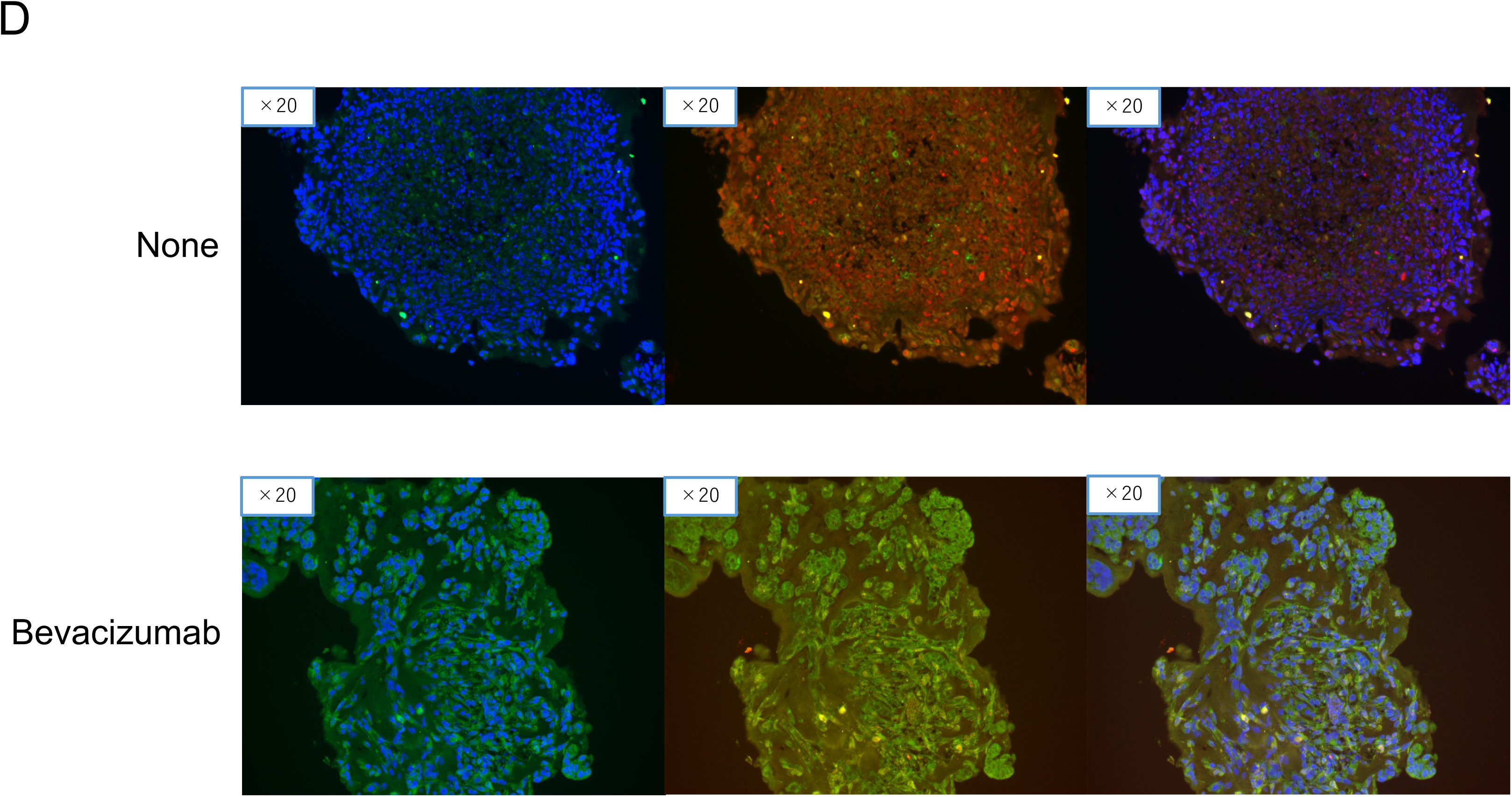
Analysis of the combined effects of Bevacizumab (Bev) and EGFR-TKI on EGFR-mutant cancer cells. (A) and (B) depict the results of a luciferase assay measuring cell viability following Osimertinib treatment alone or in combination with Bev in HCC827 and H1975 organoids, respectively (n=3, *p<0.05, Mann-Whitney U test). (C) Immunofluorescence analysis of E-cadherin and ZEB-1 in HCC827. E-cadherin (green; left panels) and ZEB-1 (red; middle panels) expression and localization in HCC827 organoids, with and without Bevacizumab treatment. Merged images are shown in the right panels. (D) Corresponding analysis in H1975 organoids.

Further examination of the EMT changes in response to Bev treatment was undertaken via immunofluorescence analyses in HCC827 organoids (Figure 4C). The results revealed an increase in E-cadherin-positive cells and a reduction in ZEB-1-positive cells, implying that the addition of Bev might attenuate EMT. A corresponding observation was made in the H1975 organoids (Figure 4D). Collectively, these results offer a promising direction for improving EGFR-TKI resistance in EGFR-mutant lung cancer. The observed EMT modulation by Bev highlights the potential utility of this VEGF inhibitor in conjunction with EGFR-TKIs for more effective treatment strategies.

### Bevacizumab Inhibits on EMT in Cancer Organoids

The possible role of Bevacizumab in the inhibition of EMT within cancer organoids was scrutinized. The expression of E-cadherin and ZEB-1, critical markers of EMT, was assessed by Western blotting following the culture of HCC827 cells (Figure 5A) and H1975 cells (Figure 5B) with Bevacizumab.

**Figure 5:**
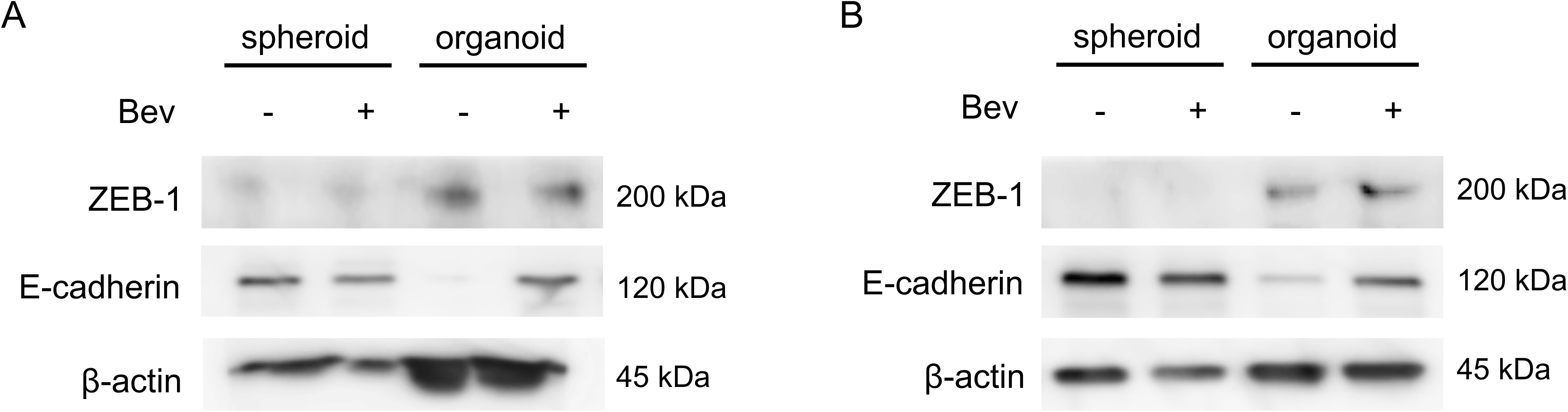
Western blot analysis examining E-cadherin and ZEB-1 expression in spheroids and organoids, with and without Bevacizumab treatment. (A) ZEB-1 (top) and E-cadherin (middle) levels in HCC827 spheroids and organoids are presented. β-actin is utilized as a loading control (bottom panel). (B) The same analysis is conducted for H1975.

The data presented in Figures 5A and 5B reveal that E-cadherin expression was enhanced in organoids treated with Bevacizumab, in contrast to those without the treatment. This upregulation of the epithelial marker E-cadherin is indicative of a suppression in EMT, thereby suggesting an ameliorative effect of Bevacizumab on the EMT process.

### miR200c Modulates EMT and Enhances EGFR-TKI Sensitivity in Organoids

As previous reports have suggested that miR200c reduces EMT in cancer cells through the regulation of ZEB-1[14], we aimed to investigate the role of miR200c in modulating the sensitivity of organoids to EGFR-TKIs. To this end, a mimic of miR200c was transfected into the organoids, and a significant increase in miR200c levels was observed in H1975 organoids compared to controls (Figure 6A).

**Figure 6:**
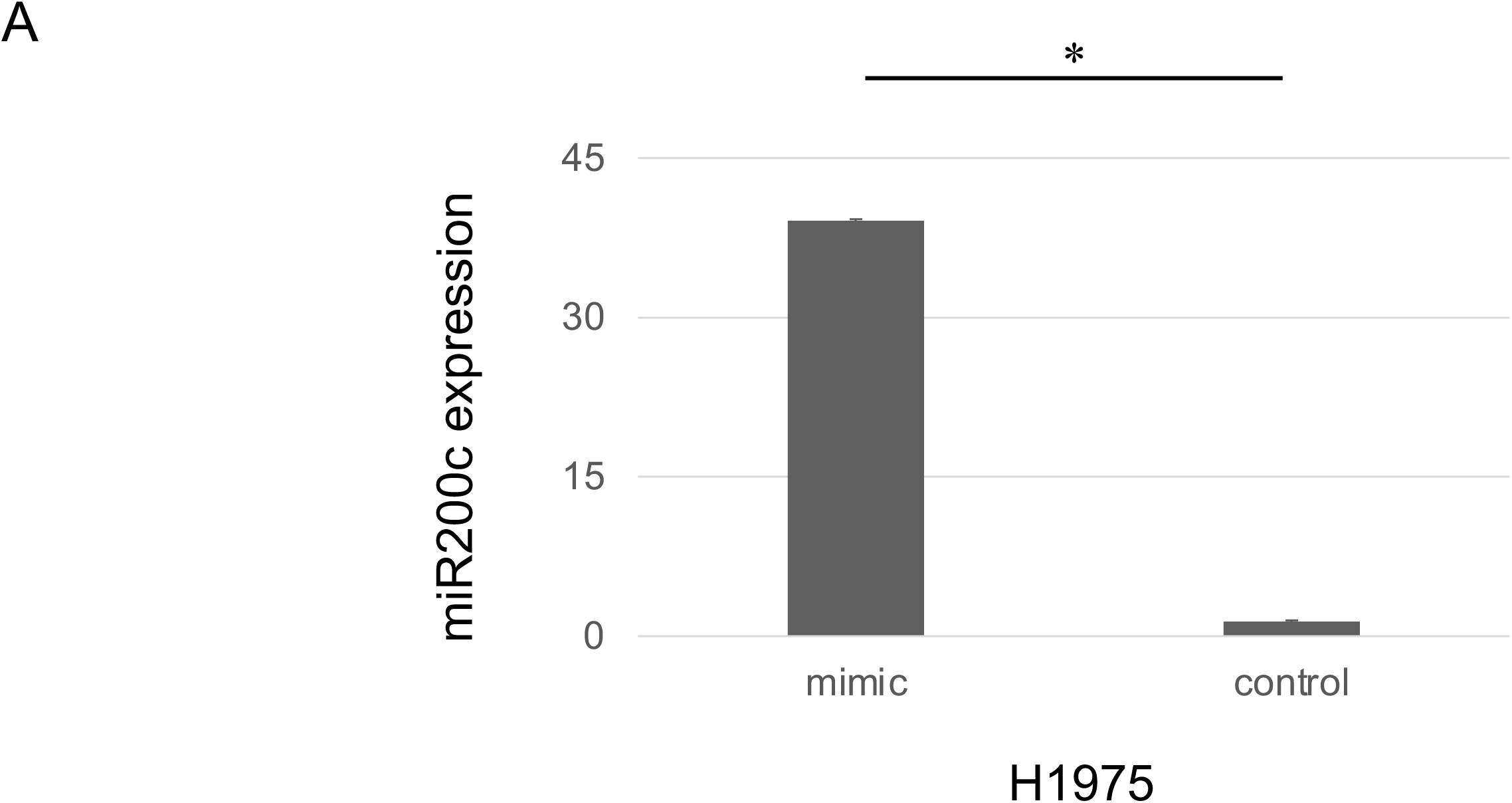
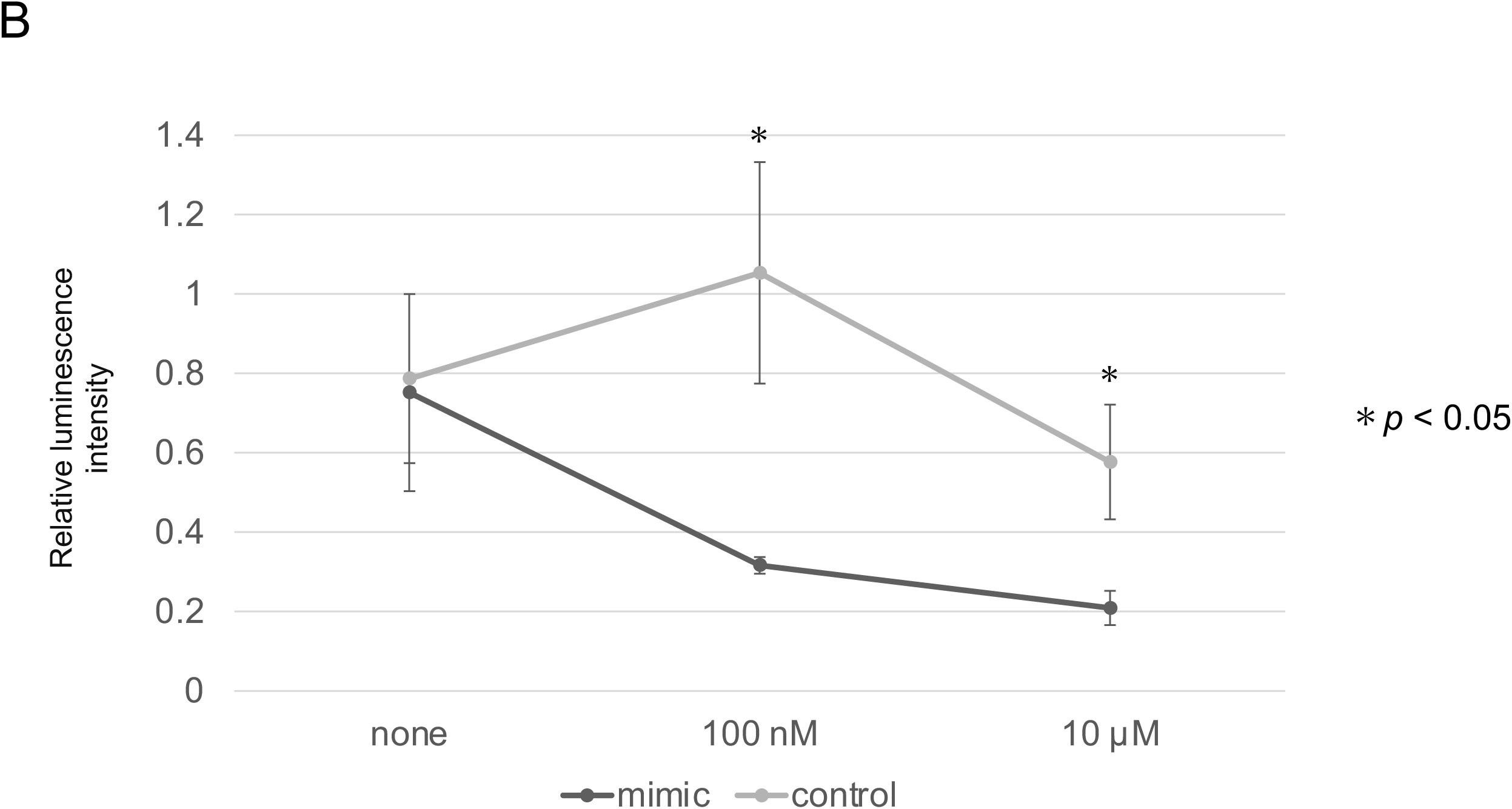
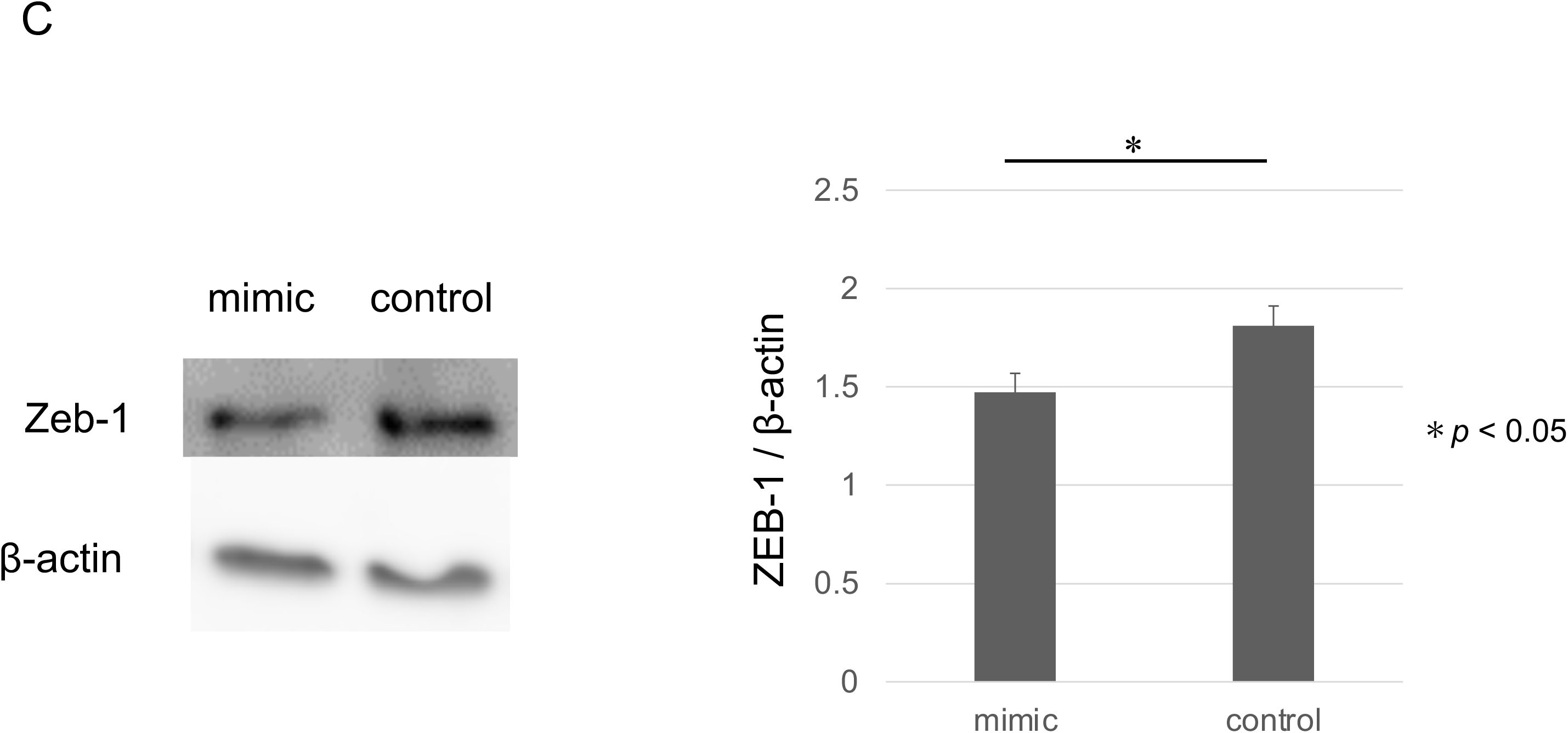
The effects of miR200c on EGFR-mutant lung cancer organoids. (A) RT-PCR analysis comparing the expression of miR200c between organoids transfected with miR200c mimic and control organoids (n=3, *p<0.05, Mann-Whitney U test). (B) Luciferase assay depicting cell viability in organoids treated with Osimertinib with or without the addition of miR200c mimic (n=3, *p<0.05, Mann-Whitney U test). (C) Western blot analysis in spheroids and organoids with or without the introduction of miR200c mimic, assessing the expression of ZEB-1 and β-actin. The bar graph, created using Image J, compares protein quantifications (n=3, *p<0.05, Mann-Whitney U test).

Following this, the impact of the miR200c mimic on the susceptibility of organoids to EGFR-TKIs was assessed via a luciferase assay. The IC50 of organoids transfected with the miR200c mimic was significantly lower than that of the controls, suggesting enhanced drug sensitivity (Figure 6B).

Regarding the expression level of ZEB-1, a known target of miR200c and an EMT marker, we observed a significant decrease in ZEB-1 expression in H1975 organoids after transfection with the miR200c mimic, compared to the controls (Figure 6C).

## Discussion

We established spheroids and organoids by culturing lung cancer cell with EGFR mutation, MSC and HUVEC on matrigel. The results of cell viability analysis indicated that the organoids had the drug resistance compared to the spheroids. The increase of ZEB-1 and the decrease of E-cadherin were shown in the organoids, which indicated that the drug resistance might be caused by EMT. Adding Bev as well as miR200c to the organoids resulted in the improvement of the drug resistance and EMT.

EGFR-TKIs have significantly improved treatment for NSCLC patients with activating EGFR mutations. However, TKI resistance often develops, leading to disease progression typically within a couple of years. Therefore, finding new drugs or strategies to counteract TKI resistance and enhance patient outcomes in advanced NSCLC is a priority[15], [16].

EMT and its role in drug resistance are emerging areas of interest in cancer research. Numerous EMT-related signaling pathways have been implicated in cancer cell drug resistance [17]. Furthermore, EMT is one of key resistance mechanisms against EGFR-TKIs including other mechanisms such as the ‘gatekeeper’ T790M mutation, histologic transformation from NSCLC to small-cell lung cancer, and concurrent molecular or genetic alterations [18]–[20]. Additional mechanisms include activation of alternative signaling pathways like MET amplification, HER2 overexpression [21], [22]. Research has capitalized on in vitro models to explore resistance mechanisms to anti-cancer drugs, specifically those linked to EMT. These models offer a controlled environment for understanding the intricacies of drug resistance and for trialing novel therapeutic approaches. Liu et al., probed the association between cancer cell stemness and EMT characteristics within drug-resistant esophageal cancer cells [23].

In this, we utilized organoid models comprising MSCs and HUVECs. This facilitated the examination of in vitro interactions between diverse cellular entities within the TME [24]. Notably, when MSCs are cultured alongside cancer cells, they can potentially give rise to cancer-associated fibroblasts (CAFs)[25]. These CAFs are critical components of the TME. Our organoids demonstrated a significant presence of stromal tissues (Figure 1), possibly representative of CAFs within the TME. Interestingly, the phenomenon of EMT can be stimulated by CAFs within tumor cells [26]. One important marker of EMT is the downregulation of E-cadherin, a protein critical for cell-cell adhesion within epithelial tissues. Notably, decreased E-cadherin levels have been correlated with EMT and poor prognoses in lung cancer patients [27], [28]. Additionally, ZEB 1, a transcription factor, is known to be inversely correlated with E-cadherin expression, further underlining its role in EMT[29]. In line with these findings, our study revealed decreased E-cadherin and increased ZEB 1 expressions within the organoids (Figures 2, 3). This shift in protein expression potentially indicates the induction of EMT and resultant drug resistance, underscoring the relevance of our model in studying these critical processes in NSCLC.

Bevacizumab, an anti-angiogenic agent, functions by inhibiting VEGF signaling pathways. Its dual effects involve impeding angiogenesis to deprive the tumor of oxygen and normalizing the vasculature to enhance treatment sensitivity[30]. The addition of Bevacizumab to Erlotinib has been observed to extend progression-free survival in NSCLC patients compared to Erlotinib monotherapy[31]. In our investigation, Bevacizumab appeared to ameliorate drug resistance and EMT (Figures 4-6). There is evidence to suggest that an elevated VEGF expression in malignant cells may drive EMT[32]. Existing literature presents conflicting findings about the relationship between Bevacizumab and EMT. For instance, Kim et al. demonstrated that Bevacizumab could inhibit TGF-β1 induced EMT in colon cancer cells[33]. However, in a stark contrast, another study reported Bevacizumab to promote EMT through Wnt/β-catenin signaling in glioblastoma cells. Further research should focus on clarifying the role of VEGF and the effects of Bevacizumab[34].

We also explored the role of miR200c, in relation to EMT in cancers organoids. Prior research has shown that miR200c plays a pivotal role in EMT among several cancers[35]. In EGFR-mutant lung cancer cell lines displaying EMT features, a downregulation of miR200c was observed, attributable to the increased level of ZEB-1 in these cells[36]. ZEB-1, an EMT-promoting transcription factor, appears to be regulated by miR200c. Upon transfecting organoids, which exhibited a more pronounced EMT than cell lines, with miR200c, we noticed an improvement in EMT and drug resistance, accompanied by a decrease in ZEB-1 (Figure 6). This underscores the potential of miR200c in future EMT-targeting strategies aimed at overcoming resistance to chemotherapy.

Despite the several novel findings, our study is not without limitations. Firstly, the TME replicated in our organoids doesn’t fully encapsulate the intricate characteristics of the TME in human cancers. Elements such as immune cells, exosomes, and vessels, which play a crucial role in the human cancer milieu, are absent in the organoids. Thus, there is an imperative need to validate these results in in vivo models. Secondly, the lung cancer cells employed in this study harbor specific EGFR mutations (del 19 or L858R/T790M), which could potentially bias the results. Clinical trials have indicated variances in the efficacy of EGFR-TKIs across different gene alterations. Hence, these findings need to be interpreted in light of these specificities.

## Conclusion

In conclusion, our study provided compelling evidence suggesting that EMT is a key mechanism of resistance against EGFR-TKIs, emphasizing the necessity of EMT-targeting strategies in overcoming this resistance. Our in vitro organoid models offered valuable insights into the tumor microenvironment, while the role of Bevacizumab and miR200c in modulating EMT was further highlighted. Future research should aim to elucidate these interactions more precisely and explore their potential in clinical applications.

## CRediT authorship contribution statement

Nobuaki Kobayashi: Conceptualization, Methodology, Validation, Formal analysis, Investigation, Data Curation, Writing - original draft, Visualization. Seigo Katakurai: Conceptualization, Methodology, Data curation, Visualization. Nobuhiko Fukuda: Methodology, Data curation, Writing - review & editing. Shuhei Teranishi: Data curation, Supervision. Sousuke Kubo: Data curation. Chisato Kamimaki: Investigation, Data curation. Hiromi Matsumoto: Supervision. Kohei Somekawa: Data curation. Ayami Kaneko: Data curation. Ishikawa Yoshihiro: Supervision. Koji Okudela: Supervision. Keisuke Sekine: Supervision. Takeshi Kaneko: Supervision. Nobuaki Kobayashi, Seigo Katakura, and Nobuhiko Fukuda contributed equally to this work. They are co-first authors of this article.

## Declaration of generative AI and AI-assisted technologies in the writing process

During the preparation of this work the author(s) used chatGPT in order to improve readability. After using this tool/service, the author(s) reviewed and edited the content as needed and take(s) full responsibility for the content of the publication.

## Abbreviations

Bev: Bevacizumab - A monoclonal antibody that can interrupt vascular growth by binding to vascular endothelial growth factor.
CAFs: Cancer-associated fibroblasts - Cells within the tumor microenvironment that promote tumorigenesis.
EGFR: Epidermal Growth Factor Receptor - A protein present on certain cell surfaces where epidermal growth factor binds, causing the cells to divide.
EMT: Epithelial-Mesenchymal Transition - A process whereby epithelial cells lose cell polarity and cell-cell adhesion, and gain migratory and invasive properties.
HUVECs: Human Umbilical Vein Endothelial Cells - Cells derived from the endothelium of veins from the umbilical cord.
MSCs: Mesenchymal Stem Cells - Multipotent stromal cells that can differentiate into a variety of cell types.
NSCLC: Non-small Cell Lung Cancer - The most common type of lung cancer.
TME: Tumor Microenvironment - The environment around a tumor, including the surrounding blood vessels, immune cells, fibroblasts, signaling molecules and the extracellular matrix.
TKIs: Tyrosine Kinase Inhibitors - A type of drug that inhibits tyrosine kinases, enzymes responsible for the activation of many proteins by signal transduction cascades.
VEGF: Vascular Endothelial Growth Factor - A signal protein produced by cells that stimulates the formation of blood vessels

